# Double Emulsion Picoreactors for High-Throughput Single-Cell Encapsulation and Phenotyping via FACS

**DOI:** 10.1101/2020.06.07.139311

**Authors:** Kara K. Brower, Margarita Khariton, Peter H. Suzuki, Chris Still, Gaeun Kim, Suzanne G. K. Calhoun, Lei S. Qi, Bo Wang, Polly M. Fordyce

**Affiliations:** Department of Bioengineering, Stanford University, Stanford, CA 94305; Chem-H Institute, Stanford University, Stanford, CA 94305; Institute for Stem Cell Biology and Regenerative Medicine, Stanford CA 94305; Department of Chemical Engineering, Stanford University, Stanford, CA 94305; Department of Chemical and Systems Biology, Stanford University, Stanford, CA 94305; Department of Developmental Biology, Stanford University, Stanford, CA 94305; Department of Genetics, Stanford University, Stanford, CA 94305; Chan Zuckerburg BioHub, San Francisco, CA 94158

**Keywords:** single cell analysis, droplets, double emulsions, FACS, microreactors, cellular phenotyping, microfluidics

## Abstract

In the past five years, droplet microfluidic techniques have unlocked new opportunities for the high-throughput genome-wide analysis of single cells, transforming our understanding of cellular diversity and function. However, the field lacks an accessible method to screen and sort droplets based on cellular phenotype upstream of genetic analysis, particularly for large and complex cells. To meet this need, we developed Dropception, a robust, easy-to-use workflow for precise single-cell encapsulation into picoliter-scale double emulsion droplets compatible with high-throughput phenotyping via fluorescence-activated cell sorting (FACS). We demonstrate the capabilities of this method by encapsulating five standardized mammalian cell lines of varying size and morphology as well as a heterogeneous cell mixture of a whole dissociated flatworm (5 - 25 μm in diameter) within highly monodisperse double emulsions (35 μm in diameter). We optimize for preferential encapsulation of single cells with extremely low multiple-cell loading events (<2% of cell-containing droplets), thereby allowing direct linkage of cellular phenotype to genotype. Across all cell lines, cell loading efficiency approaches the theoretical limit with no observable bias by cell size. FACS measurements reveal the ability to discriminate empty droplets from those containing cells with good agreement to single-cell occupancies quantified via microscopy, establishing robust droplet screening at single-cell resolution. High-throughput FACS phenotyping of cellular picoreactors has the potential to shift the landscape of single-cell droplet microfluidics by expanding the repertoire of current nucleic acid droplet assays to include functional screening.

**ABSTRACT FIGURE:** 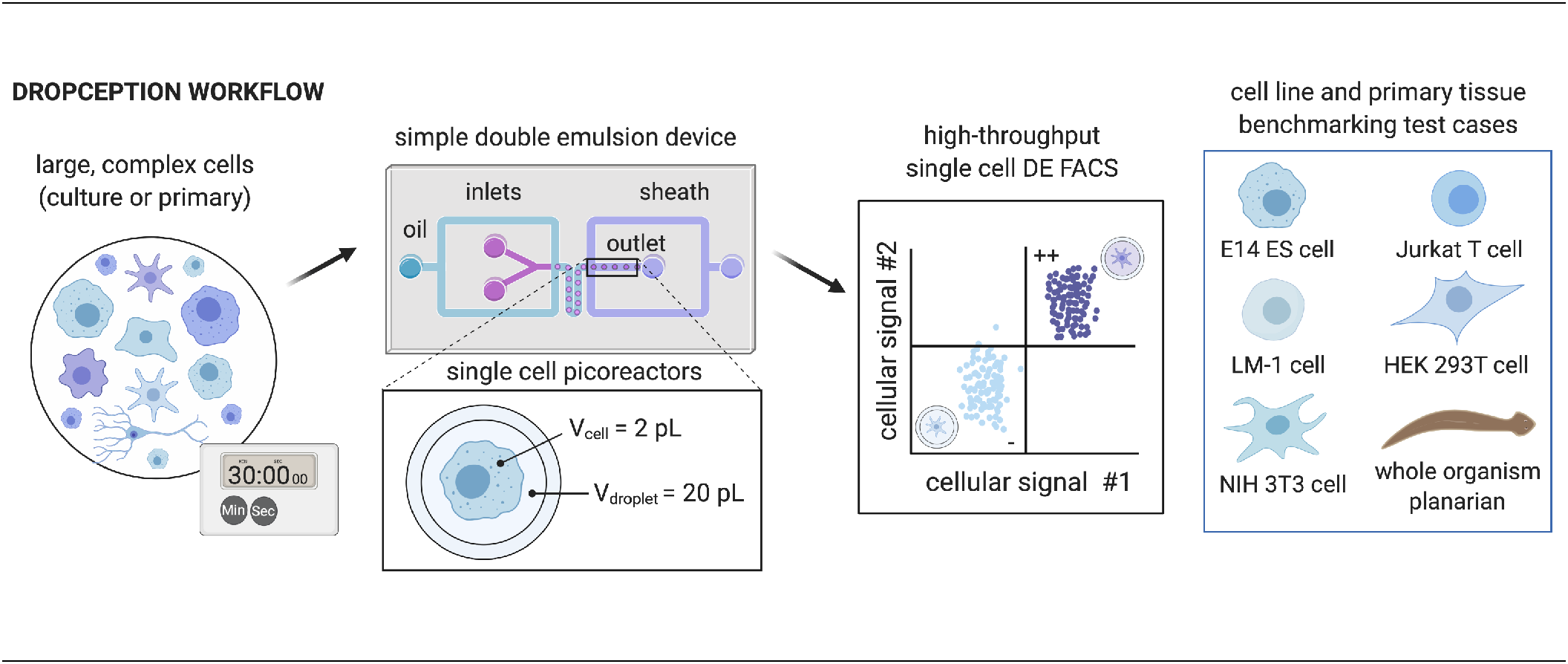

## INTRODUCTION

The last decade has yielded an exponential rise in new methods to analyze single cells^1,2^, revealing critical insights into cellular diversity^3–7^, tissue organization^8–10^, and organism development^11–14^. In particular, droplet microfluidics has emerged as a powerful class of single-cell isolation techniques due to its unprecedented scale (0.1 - 10M droplets per run), throughput (0.1 - 30 kHz generation rate), and efficiency ($0.10 - $0.50 per cell)^15–17^. Novel droplet assays have enabled thousands of single cells to be profiled by genome^18,19^, epigenome^20,21^, transcriptome^3,4,22^, or proteome^23–25^, leading to the generation of the first whole-organism cell atlases^26–29^. Due to their ease-of-operation and low barrier to entry, open-source droplet technologies (e.g., DropSeq^3^, InDrops^4^) have been adopted by specialists and non-specialists alike with commercial droplet platforms (e.g., 10X Genomics^20,22^) achieving wide-spread market penetration in research laboratories world-wide^15,30^.

Despite these advances, single-cell droplet techniques remain fundamentally limited in their ability to easily screen droplets based on phenotypic signals^16^. This capability would enable new opportunities for single-cell analysis. First, isolating and sequencing only those droplets containing cells would dramatically lower assay costs^15^ while increasing sequencing accuracy and depth^31–33^. Second, encapsulated cells could be isolated based on phenotypes not currently measurable with standard fluorescence-activated cell sorting^34,35^ (FACS), such as enzymatic turnover, presence of secreted molecules, or quantification of proteins lacking cell surface markers^17^. Lastly, while droplets have been used to perform either single-cell phenotyping^36–40^ (*e.g.,* secreted marker screens, metabolite profiling, enzyme kinetic assays) or genome-wide sequencing^3,4,18,20–22^ (*e.g.,* RNA-seq, ATAC-seq, WGA analysis), no technique combines the two for multi-omic measurements. Sorting individual droplets by their cellular biochemical signals with downstream genome-wide profiling of the same cell would directly link cellular phenotypes to their underlying genetic mechanism^3,4,16^.

Most single-cell droplet assays employ water-in-oil (W/O) single emulsions in which a large (~1 nL) aqueous droplet containing cells and reagents is surrounded by oil^15^. However, sorting these droplets can only be done via fluorescence-activated droplet sorting (FADS)^41–43^, which requires extensive instrumentation, custom optics, and technical expertise to build and operate, severely limiting the applicability of the technique. As a result, there exists no easily accessible means to screen and sort droplets by cellular presence or functional perturbation response, preventing translation of powerful droplet-based sequencing technologies^3,4,20,22^ to phenotypic multi-omic profiling.

A promising alternative involves encapsulating cells within double emulsions (DE) (water-oil-water, W/O/W)^39,40,44^. Unlike W/O droplets, DE droplets can be suspended in aqueous solutions, making them compatible with standard FACS instruments^45^. DEs have previously been combined with FACS for screening of bacterial or yeast mutant libraries^39,46,47^. However, these techniques suffer unpredictable cell occupancy and high multiple-cell loading (‘multiplet’) rates, confounding the downstream phenotype to genotype linkage. No prior effort has demonstrated successful encapsulation of large animal cells within picoliter-scale DE droplets, likely due to challenges associated with encapsulating large cells within droplets that are sufficiently small to pass through FACS nozzles without breakage and cross-contamination^45^.

Recently, we developed a new method^48^ (sdDE-FACS, for <u>s</u>ingle <u>d</u>roplet <u>d</u>ouble <u>e</u>mulsion FACS) to sort and recover large DE droplets via FACS by internal droplet fluorescence signals with similar performance to single-cell FACS (>70% sort recovery, 99% target sensitivity) using commercially available cytometers. Using sdDE-FACS, we established the first reliable isolation of single droplets based on florescence phenotype and recovered encapsulated nucleic acids at high efficiency post-sort.

Building on this progress, we present here the first demonstration of high-throughput FACS-based screening of picoliter-scale droplets containing single animal cells. Using a custom microfluidic device, we demonstrate a simple workflow, Dropception, for encapsulating large, complex cells (5-25 μm in diameter) within highly monodisperse DE droplets small enough (~45 μm in total diameter) for FACS. We precisely tuned droplet size and cell concentration for an extremely high ratio of single- to multiple-cell loading events^49^. We benchmark performance of this technique across 5 standard mouse and human cell lines (**Table 1)** for robust encapsulation of single cells near maximal theoretical loading efficiency with no observable cell size bias. Using a modified sdDE-FACS workflow for large droplets, we screen tens of thousands of cell-containing DEs within minutes via a standard flow cytometer, establishing accurate discrimination of single-cell droplets from empty droplets. Finally, we apply Dropception to heterogeneous cell populations collected from a whole flatworm planarian, illustrating the wide applicability of this technique to a variety of different cell types and primary samples.

**Table 1.** Standard cell lines used for performance characterization in this study.

| Cell line | Description | Mean diameter (μm) | Source |
| --- | --- | --- | --- |
| **HEK 293T** | Human embryonic kidney cell | 13.7 ± 2.2 | Takara Bio 632180 |
| **Jurkat, Clone E6-1** | Human T lymphocyte | 12.0 ± 2.0 | ATCC TIB-152 |
| **E14** | Mouse embryonic stem cell | 11.7 ± 1.2 | ATCC CRL-1821 |
| **NIH 3T3** | Mouse embryonic fibroblast | 17.8 ± 2.6 | Sigma 93061524 |
| **LM-1** | Mouse macrophage cell line | 14.5 ± 2.7 | Kerafast EMG014 |

## RESULTS AND DISCUSSION

### THE DROPCEPTION WORKFLOW

The Dropception workflow takes place in two stages (**Fig. 1**): (1) encapsulation, in which cells and reagents are introduced into a one-step microfluidic device to yield a library of uniform, picoliter-scale DE droplets, and (2) screening, in which these DE droplets are passed through a FACS machine for high-throughput analysis by cellular presence or phenotype. To facilitate adoption of the technique, we employ a widely available commercial flow cytometer and our droplet generation device requires only 4 syringe pumps and an inexpensive benchtop microscope for operation (**Table S1**).

**FIG 1.**
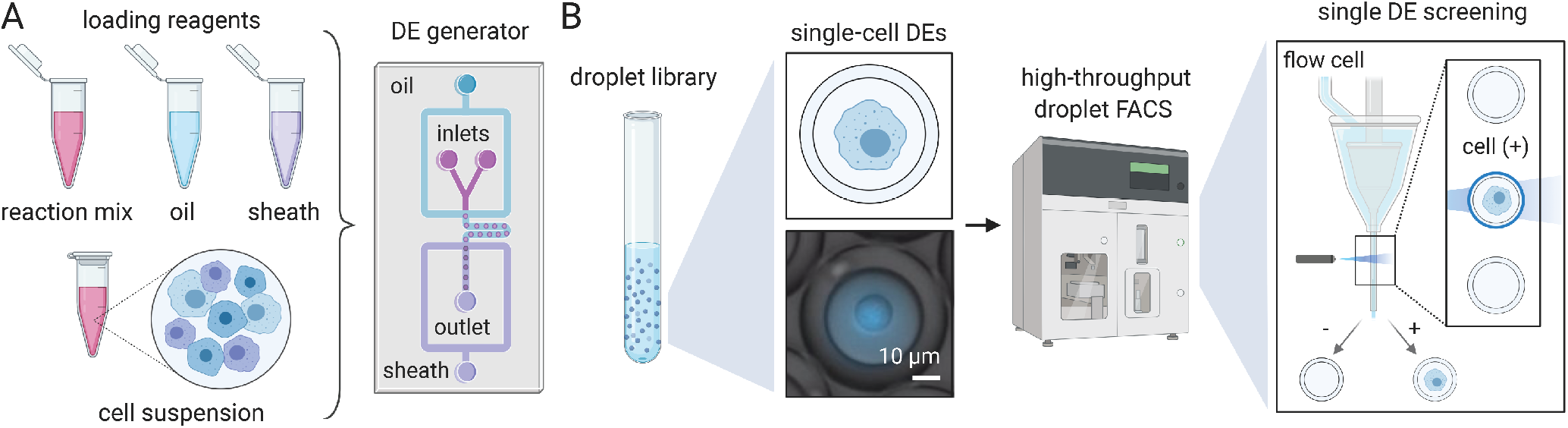
Schematic illustration of the Dropception workflow for cell encapsulation and droplet phenotyping. **(A)** DE picoreactors are generated from single-cell suspension and co-encapsulated with a reagent mixture in an oil shell and surrounding sheath buffer. **(B)** Droplet FACS is conducted on standard commercial flow cytometers at 12-14kHz using protocols optimized for large droplet stability and recovery. Droplet generation takes 30 min and FACS phenotyping of >50,000 droplets takes 5 min. The microscopy image shows a mouse ES cell encapsulated in a DE droplet (Calcein AM, blue).

This workflow addresses several technical challenges required for high-throughput screening of cell-containing picoliter-scale droplets (‘picoreactors’). First, double emulsions must be small enough for FACS yet large enough to encapsulate mammalian cells reliably^50^ and must remain stable during droplet generation and flow cytometry^48,51,52^. Second, FACS must be able to accurately discriminate between cell-containing droplets and empty droplets and, ideally, associate fluorescence signals with encapsulated *single* cells^3,4,22^. Lastly, the workflow must be compatible with multiple cell types and in-droplet reaction schemes to facilitate translation and broad applications^16^.

To address these challenges, we designed a custom microfluidic device for large cell encapsulation into picoliter double emulsions capable of FACS analysis and sorting. By generating uniform droplets at picoliter scale via a specific loading distribution (Poisson, λ<0.05), we ensure cell-containing droplets achieve high single-cell purity (>98% of cell-containing droplets are single cells) without compromising low reagent consumption, a common pitfall of large-droplet techniques^49^. Our workflow enables a variety of potential reaction schemes; picoliter droplet reactions using our one-step device can co-encapsulate lysis and reaction solutions for genomic and transcriptomic profiling, secreted marker analysis, or enzymatic turnover. Each experiment takes less than 30 min including cell staining, minimizing changes to the native state of encapsulated cells^53^ (**Figs. S1, S2**).

### DEVICE DESIGN AND CHARACTERIZATION

High data quality in single-cell analyses depends on the ability to discern which droplets contain single cells^31–33^. Previously, it has been difficult to attain predictable single-cell loading in DEs due to droplet polydispersity^39,47^. To achieve single-cell droplet FACS, picoliter DEs needed for FACS analysis must be highly uniform in size to yield accurate cell occupancy distributions^49^. However, monodisperse DE generation is technically challenging, especially when attempting to load large particles into small droplets^54,55^. To enable robust large cell encapsulation in double emulsions, we designed a novel device containing optimized design elements for flow stability.

The Dropception device employs a dual flow-focusing geometry^56,57^ for co-encapsulation of cells and assay reagents into picoliter-scale droplets (**Fig 2A, Supplementary Information**). In the first flow focuser (FF1), cells and reagents from the inlet tree meet a stream of carrier oil and are encapsulated into regularly-spaced W/O single emulsions. In the second flow focuser (FF2), the cell-laden single emulsions in their carrier oil meet an aqueous stream and are pinched off to form W/O/W double emulsion droplets, each containing an oil shell and aqueous interior. The carrier oil is a biocompatible fluorocarbon oil optimized for high oxygen delivery to encapsulated cells^58,59^. Device operation requires only 100 μL of cell suspension or reaction mix (compared to >1000 μL in techniques such as DropSeq^3^) with minimal reagent consumption per droplet picoreactor, enabling screening of precious samples.

**FIG. 2.**
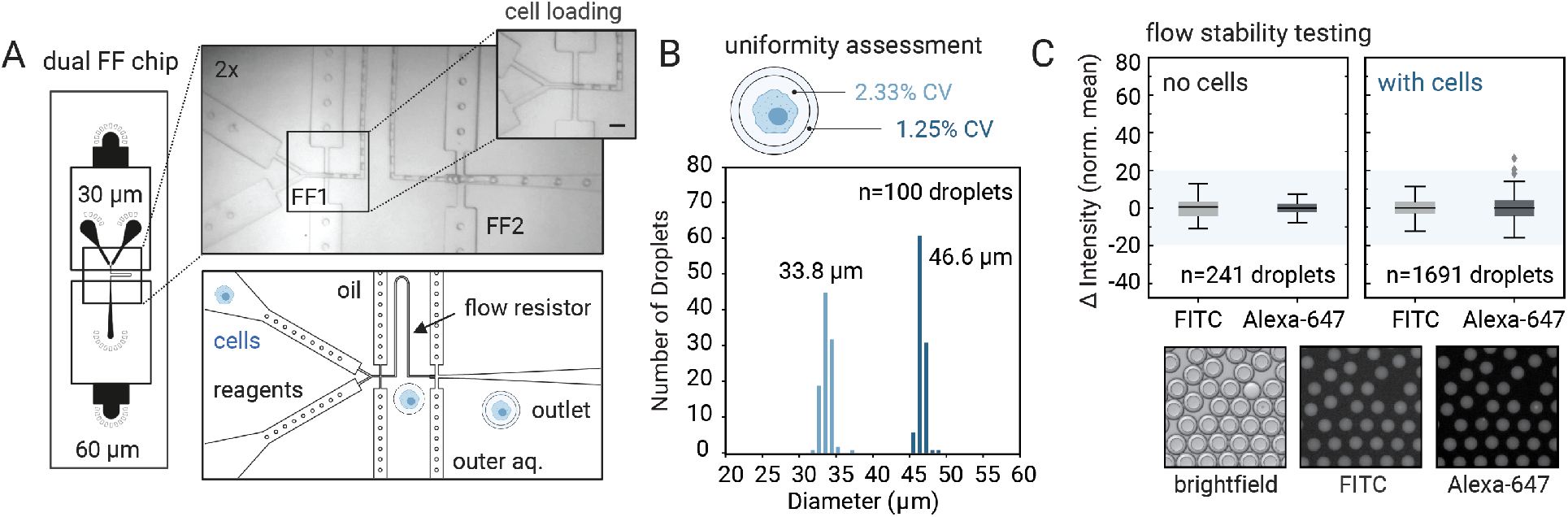
The Dropception device generates monodisperse droplets under stable flow. **(A)** Design and microscopy image of the device showing flow focuser features (FF1, FF2), inlets, channels, and outlet. *Inset*: cell loading at the inlet tree; flow line delineates relative volumetric contributions of inlets. Scale bar: 45 μm. **(B)** DE size characterization via microscopy denoting internal core (*light blue*) and total droplet (*dark blue*) diameters with corresponding coefficients of variation (CV) (n=100, sample includes cells). **(C)** Two-dye co-flow experiments with and without cells show flow stability across droplet populations; intensity is normalized to zero-mean (interquartile ranges: (−2.16, 2.68) and (−2.32, 2.21) for FITC; (−2.20, 1.95) and (−2.70, 2.82) for Alexa-647 in the absence and presence of cells, respectively).

Upstream of the first flow focuser, we designed an inlet tree containing two wide channels without flow filters, each spaced 30° to normal, which funnel into short resistive elements to focus flow at a short channel (**Fig 2A**). We included a short flow resistor and short resistive elements at each flow focuser to produce ordered, triggered flow^60,61^ where each aqueous single emulsion is encased in an oil emulsion to create a double emulsion at efficiencies beyond stochastic statistics (>99.9% of droplets contain a single emulsion core).

To minimize cross-contamination between droplets, the cell and reagent inlet channels meet just 110 μm before the FF1 nozzle (below Peclet diffusion distance). During operation, cells are suspended in a density gradient medium (20% OptiPrep) to avoid settling in the loading syringe. In this region, the difference between the index of refraction of the cell solution and the reaction mix allows for clear delineation of the relative contributions of each inlet (inset, **Fig. 2A**), thereby serving as a precise visual readout to tune relative reaction volumes in the droplet core during operation^62^ (**Supplemental Methods**).

The Dropception device has channel heights and nozzle widths of 30 and 22.5 μm for the FF1 region and 60 and 45 μm for the FF2 region. Droplet generation with cell encapsulation produced DEs with diameters of 34 and 47 μm for the inner aqueous core and outer oil shell, respectively (20 and 54 pL by volume) (representative population, **Fig 2B)**. Matching the flow rate of the outer aqueous sheath to the periodicity of single emulsion generation yielded highly monodisperse DE populations with uniform inner aqueous core sizes (2.33% and 1.25% CV on the inner core and outer shell diameters, respectively; **Fig. 2B**). These results were consistent across all subsequent data presented here (**Figs. 3, 4, 5, S1**) using a single set of flow conditions (**Table S2**).

**FIG. 3.**
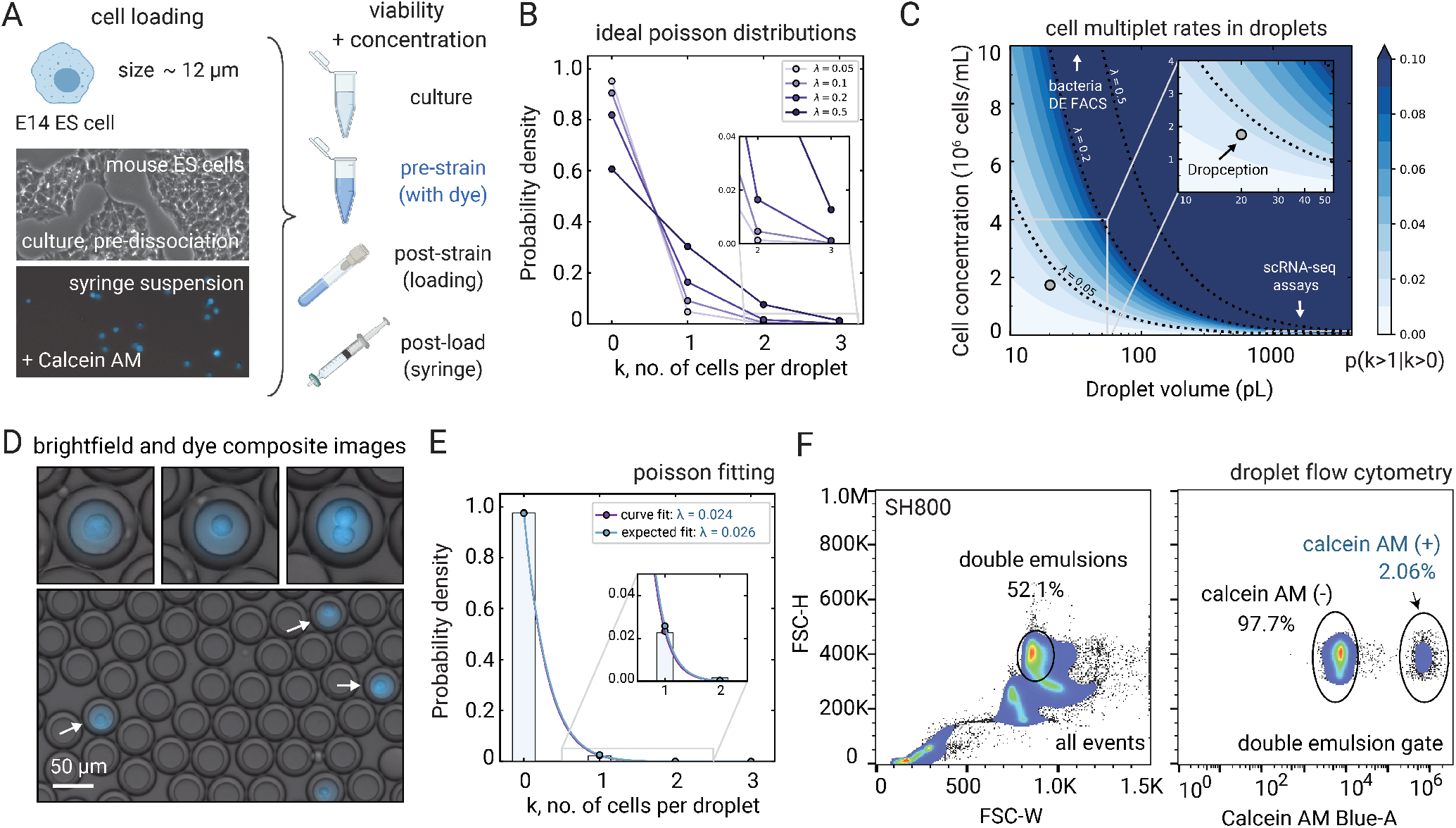
ES cell encapsulation approaches theoretical loading with single-cell DE discrimination by FACS. **(A)** Microscopy images show mouse E14 cells from culture and syringe suspension during processing, with cartoons depicting additional concentration and viability measurement steps (**Table 1**). **(B)** Expected droplet occupancy distributions across modeled Poisson loading regimes by event rate (*λ*). **(C)** Multiplet loading probabilities (as a percentage of cell-containing droplets) under typical Poisson droplet encapsulation varying by cell concentration and droplet size, with arrows highlighting comparable technologies. Inset shows the low Poisson regime chosen for Dropception. **(D)** Microscopy images of loaded ES cells in DE droplets. Arrows indicate single-cell loading (*k=2* inset shows a rare multiplet event). **(E)** Microscopy-determined cell occupancy fitted to a Poisson and plotted against expected distribution. Inset shows singlet versus doublet cell occupancy (n=7,104 droplets). **(F)** FACS analysis of DE droplets containing ES cells. DEs were gated on forward scatter across all events (FSC-H vs. FSC-W, *left*) and analyzed under a violet laser for Calcein AM (right). DE-gated population (*right*) shows two clearly separable populations (cell containing (+) *vs* empty (−) droplets (n=45,000 droplets).

**FIG. 4.**
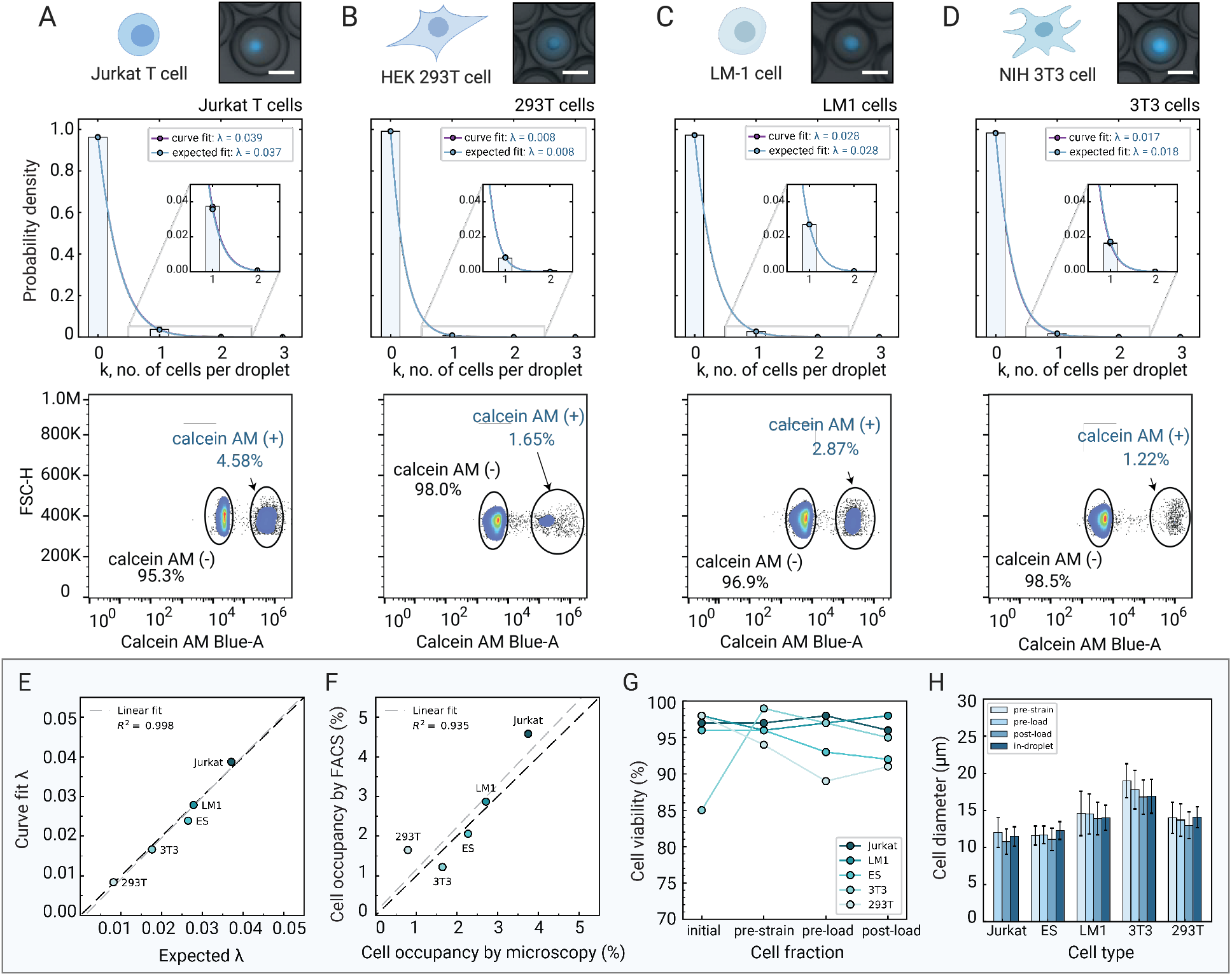
Benchmarking single-cell encapsulation and phenotyping of Jurkat T, HEK 293T, LM-1, and 3T3 cell lines in DE droplets. **(A, B, C, D)** Microscopy images (*top*), cell occupancy distributions (*middle*) (n=4,028-7,449 droplets) and FACS phenotyping (*bottom*) (n=45,000 droplets) of the 4 cell lines in DE picoreactors. Scale bars: 25 μm. **(E)** Fitted versus expected event parameter for Poisson loading across all cell lines**. (F)** Cell occupancy determined by FACS plotted against single-cell occupancy determined by microscopy counts. Dashed gray lines depict linear curve fits. **(G)** Cell viability measurements during processing steps in the cell encapsulation workflow, including suspension post-loading. **(H)** Cell diameters as measured in processing fractions or within the droplet volume during the workflow across all cell lines.

**FIG. 5.**
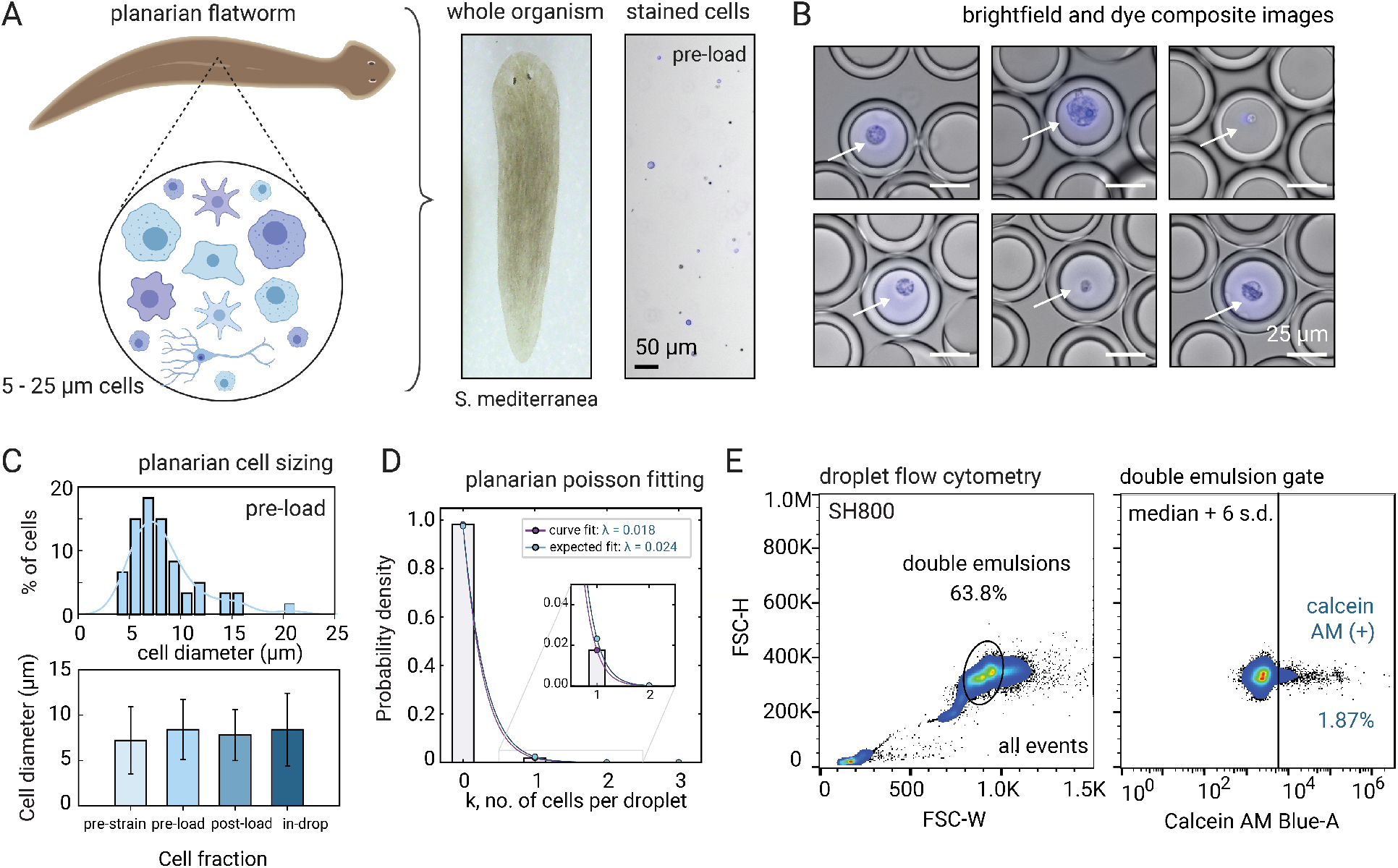
Single-cell DE encapsulation of dissociated planarian cells suggests robust applicability to heterogeneous primary tissue. **(A)** Illustration and microscopy images of complex cell populations in the planarian (Calcein AM, blue). **(B)** Representative microscopy images of *S. mediterranea* cells in DE droplets highlighting size variance of the encapsulated population. **(C)** Cell size distributions across workflow steps show broad variation in the pre-loaded cell fraction (*top*), with similar variance observed in droplet-loaded cells (*bottom*). **(D)** Microscopy-derived cell occupancy with associated Poisson fits. **(E)** FACS screening of DEs encapsulating planarian cells (n=43,238 droplets). Cell-positive gate was established via comparison to empty droplet populations (median + 6 s.d., Calcein AM signal).

To assess the uniformity of flow from each inlet during device operation, we introduced FITC- and Alexa-647-conjugated BSA into the cell and reagent inlets, respectively, and compared the variance in dye intensity distributions in the presence and absence of cells (**Fig. 2C**). Alexa-647 and FITC fluorescence intensity distributions associated with each droplet were narrow in both cell and cell-free conditions (**Fig. 2C**), demonstrating steady, non-pulsatile flow from each inlet. Combined, these data indicate robust operation of the Dropception device for stable droplet generation, even under large cell loading in highly constrained channels.

### SINGLE-CELL ENCAPSULATION OF MOUSE EMBRYONIC STEM CELLS

As a first application of the Dropception device and workflow, we encapsulated mouse embryonic stem (ES) cells (ATCC: E14 cell line) (**Fig. 3A**). ES cells are a critical cell line for investigating pluripotency and stemness^63^, profiling transcriptional and epigenomic reprogramming^64^, and are a vector for synthetic biology studies^65^. Mouse ES cells are also relatively small (11-13 μm diameter, 1.5 pL volume) with uniform morphology, providing a convenient experimental system to test cell encapsulation efficiency in picoliter-scale droplets (**Fig. S3**). Cells were dissociated from culture, stained with Calcein AM vitality dye, resuspended to a concentration of ~ 2.0×10^6^ cells/mL with 20% OptiPrep, and co-loaded into the device with a 0.5% BSA-PBS solution using relatively low flow rates to minimize shear stress on cells (**Video S1**). At multiple points throughout the experimental workflow, we quantified cell concentrations and assessed viability (**Table 2**).

**Table 2.**
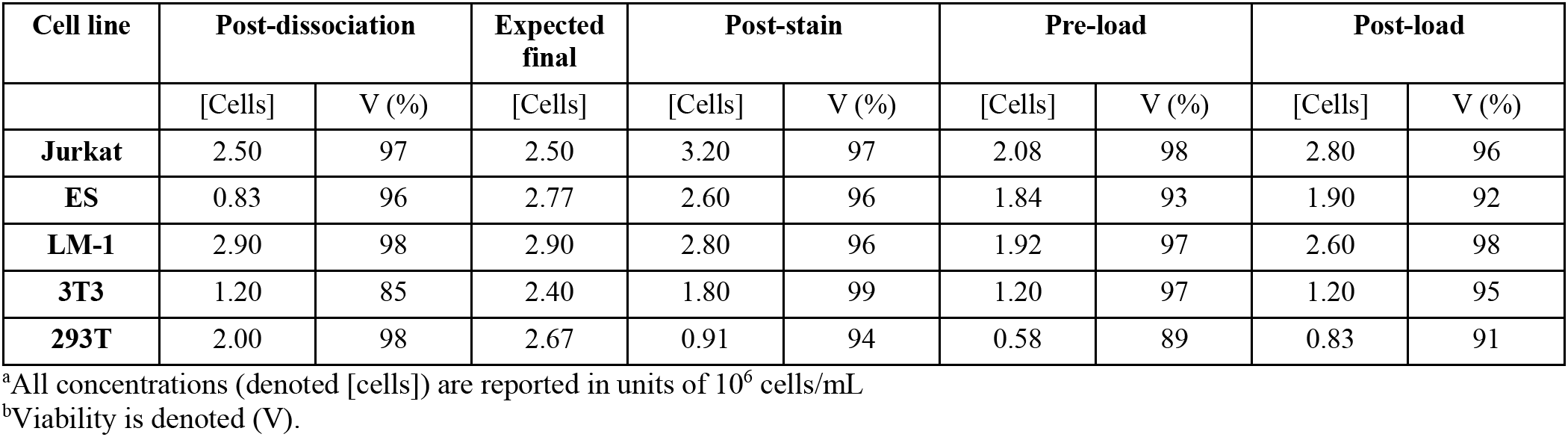
Cell viability^a^ and concentration^b^ statistics measured during the processing workflow.

Single-cell droplet encapsulation should follow a typical Poisson distribution for stochastic loading^49^ (**Fig. 3B**):

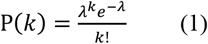

where *k* is the number of cells within each droplet and *λ* (mean number of cells per droplet) is the event rate, as given by:

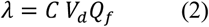

where C is the loading concentration of cells (cells/pL), V_d_ is the volume of the droplet (pL), and Q_f_ is the fraction of volumetric flow contributed by the cell suspension.

For most single-cell droplet techniques, the large majority of droplets are empty with only a small proportion containing one or more cells. Many single-cell droplet techniques optimize for single-cell occupancy (P(*k=1*)) over empty droplets (P(*k=0*)), at the expense of a high proportion of droplets containing 2 or more cells (**Fig. 3B,C)**. This fractional occupancy can be controlled by selection of droplet size and loading concentration of cells (**Fig. S4A**). Prior techniques using picoliter DEs which encapsulate yeast or bacteria at high concentration estimate up to 50% of droplets contain single cells (λ > 0.5)^39,47^. However, these techniques suffer extremely high multiplet rates (>10%), preventing true single-cell resolution in downstream sequencing.

While some mammalian cell techniques (*e.g.,* DropSeq^3^, In-drops^4^, 10X^22^) reduce multiple-cell loading events for sequencing accuracy, extremely low cell loading concentrations are required (**Fig. 3C**) and therefore reagent consumption (*e.g.,* reverse transcription enzymes, reaction components) is high to balance volumetric demands of the large droplet size. As a result, these widely-adopted sequencing technologies strike a compromise between increasing overall costs to achieve single-cell purity and reducing data quality by tolerating multiplets (~4-6% multiplet rate; inset, **Fig. S4B**).

Here, we chose to perform single-cell encapsulation under a Poisson distribution with a very low event rate (λ<0.05) via explicit selection of droplet size (~35 μm, aqueous core) and cell loading concentrations (0.5-2.5 ×10^6^ cells/mL). Cell occupancies are expected to be between 1-5% under this distribution (**Fig. 3B**). Of droplets containing cells, a minimum of 98% should contain only a single cell. Thus, this operating regime allows higher single-cell purity in the cell-containing droplet population while simultaneously significantly reducing reagent consumption via small droplet volumes (**Fig. 3C**).

To assess cell encapsulation efficiency using our workflow, we loaded ES cells into double emulsions (**Fig. 3D**). We performed manual microscopy counts of cellular occupancy across thousands of droplets and fit these data to a Poisson distribution (curve fit, **Fig. 3E**). Given known cell concentrations and droplet volume, we compared our data to ideal Poisson loading (expected distribution, **Fig. 3E**). Single-cell occupancies between the predicted and measured distributions agreed well (P(*k =* 1); 2.27% measured, 2.58% expected), with closely corresponding event rates within the intended low λ operating regime (λ: 0.024 curve fit; 0.026 expected), establishing optimal performance of cell loading near theoretical maximal loading efficiency.

### SINGLE-CELL PICOREACTOR PHENOTYPING VIA FACS

Next, we investigated whether we could discriminate cell-containing droplets from empty droplets via high-throughput DE FACS. Using a Sony SH800S FACS instrument equipped with a 130 μm sort nozzle and instrument settings from our sdDE-FACS pipeline^48^ (**Table S3**), we analyzed tens of thousands of cell-laden DE droplets (**Fig. 3F**). FACS analysis was conducted at 12 kHz with droplet sorting rates maintained below 1000 eps to achieve single droplet sort purity similar to single-cell FACS^45^. DEs comprised >50% of recorded events under low scatter thresholds that show small dust, free oil, and other debris, demonstrating high sample integrity with little evidence of droplet breakage. To the best of our knowledge, this represents the first time that droplets this large (45-48 μm) have been analyzed via FACS.

To assess whether cells encapsulated within DEs could be reliably detected via their Calcein AM fluorescence signals, we gated DE events based on their forward scatter signals (FSC-H *vs*. FSC-W, typical of large particle analysis^45^) and examined the fluorescence intensities of the gated population. Analysis of the DE gated population revealed two clearly separable populations with ~100-fold higher intensities associated with cell-containing droplets (median population intensities of 5.49×10^3^ *vs*. 4.43×10^5^). Across the total population of droplets, 2.06% were identified as containing cells (**Fig. 3F**).

This conservative FACS estimate of cellular occupancy agrees well with the microscopy-derived single-cell occupancy of 2.27%. Under the fitted Poisson distribution from empirical cell occupancies (λ = 0.024), 98.8% of cell-containing droplets should be single cells, with 1.2% remaining as multiplets in this sample. Combined, these findings suggest that picoliter DE encapsulation in a low Poisson regime (λ < 0.05) allows for high-throughput methods of analysis such as FACS to accurately assign phenotypes during droplet screening.

### SYSTEMATIC BENCHMARKING ACROSS 4 STANDARD CELL LINES

Next, we probed the experimental limits of Dropception by testing the ability to encapsulate and phenotype 4 additional standard cell lines with a wide range of morphologies and sizes (5– 20 μm) (**Table 1, Fig. S3**): human T lymphocytes and mouse macrophage cells (Jurkat and LM-1, respectively; key model systems for immunological studies^66,67^), human embryonic kidney cells (HEK 293T, common vectors for synthetic biology^68^), and mouse embryonic fibroblasts (NIH 3T3, an important resource across cancer studies^69^). All cells were loaded at a projected concentration of ~2.5×10^6^ cells/mL from dissociation (**Table 2**). Cell concentration losses were only observed for larger cell lines after straining. Similar to ES cell loading, we systematically quantified single-cell DE droplet occupancies using both microscopy and FACS.

Microscopy established successful encapsulation of all 4 cell lines (**Fig. 4**). As before, cellular occupancies (4,000 droplets/condition) were well-fit by a Poisson distribution and in agreement with expectations given observed droplet size and loading concentration for all 4 cell lines (R^2^=0.998) (**Fig. 4A-D**). This agreement demonstrates consistent performance approaching theoretical loading efficiency, even for large and morphologically diverse cell types (*e.g.* 293T and 3T3), and suggests an absence of common experimental pitfalls to cell loading such as cell clumping, flow instability, or steric bias. We found that the same flow rates and conditions could be used for efficient loading of all cell lines, suggesting that this pipeline may be used without manual adjustment for a variety of different samples and enhancing overall translatability (**Table S2, Fig. S1**).

Upon screening each of the 4 DE-loaded cell lines via FACS (**Fig. 4**), we observed clearly separable Calcein AM fluorescence intensities for gated DE populations, corresponding to empty and cell-containing droplets. The separation between these populations varied 10- to 100-fold in fluorescence intensity, indicating sufficient dynamic range to phenotype droplets by internal fluorescent signal. For all cell types, the percentage of cell-containing droplets recorded with FACS showed excellent agreement with single-cell occupancy determined by manual microscopy inspection (*R*^2^=0.935, **Fig. 4F**). Across all samples, we observed only a minor discrepancy in estimated loading rates (0.79% *vs.* 1.65% for microscopy *vs.* FACS) for HEK293T cells, likely because their relatively large size caused them to settle during FACS loading.

Finally, we characterized additional metrics of workflow performance including viability (**Figs. 4G,H**) and cell size variation (**Fig. 4H**) at all stages pre- and post-load. For all cell lines, viability remained >85% over the entire 30-minute processing window, with expected small increases in viability after filtering and pelleting to remove dead cells and debris. High cell viability during loading results in fewer free-floating nucleic acids and debris from cellular death, minimizing single-cell picroreactor cross-contamination. We observed no significant cell size distribution changes upon droplet loading even in large cell types (**Fig. 4H**), indicating an absence of steric constraint for large cell (1-3 pL) loading into 20 pL DEs.

### APPLICATION TO A WHOLE FLATWORM

Single-cell encapsulation in droplet microfluidics has largely been demonstrated using cultured cell lines, which are uniform in cell size and morphology^2,70^. However, a critical question in the field is whether new microfluidic strategies are compatible with heterogeneous cell populations freshly isolated from whole animals or tissue dissection.

To evaluate the applicability of Dropception for primary tissue, we encapsulated single cells dissociated from whole planarian flatworms (**Fig. 5A**). Planarians contain a large pool of pluripotent stem cells (neoblasts) and possess exceptional and unremitting regenerative capacity throughout their entire body, providing a powerful model organism for *in vivo* stem cell biological studies^71–75^. However, planarians are not amenable to most cell probing assays, such as transgenic marker integration or cell surface antibody markers^76^, and thus non-model organism research would benefit immensely from the ability to FACS-isolate cells by droplet-accessible phenotype (e.g., genomic PCR and secretory protein analysis)^74,75^. Furthermore, planarians have a sticky outer mucosal layer and comprise hundreds of unique cell types that vary widely in size (5-25 μm), morphology, and cellular biology^29^, posing a challenge for droplet encapsulation.

Following the same workflow as for cultured cells, we performed a fresh dissociation of whole planarians, stained cells with Calcein AM dye, suspended to a concentration of 1.63×10^6^ cells/mL with 20% OptiPrep, and loaded the resultant cell suspension into the Dropception device (**Fig. 5**). Despite the anticipated challenges, we achieved robust single-cell encapsulation of diverse cell sizes and morphologies (**Fig. 5B**). Cell size variance of dissociated primary cells was recapitulated in loaded droplets with a wide range of sizes represented in all fractions (**Fig. 5C**). Importantly, droplet counts obtained from microscopy revealed planarian single-cell occupancy in DE picoreactors was near theoretical limits (1.77%) (**Fig. 5D**).

FACS DE analysis revealed clearly discernable populations of empty and cell-containing droplets with 1.87% cell loading, in agreement with microscopy (**Fig. 5E**). Flow cytometry gates for this sample were placed conservatively by comparison to negative droplet populations (median fluorescence intensity of the Calcein AM channel (MFI: 2302) + 6 s.d. (robust s.d.: 551), consistent with empty droplet gates of cell line experiments). Broad fluorescence signals are likely due to variable dye uptake in primary cells, as only ~60% of planarian cells exhibit bright Calcein AM staining (**Fig. S5**). Under the theoretical cell encapsulation distribution for this sample (λ = 0.024), single cells should account for 98.8% of all cell-containing droplets, suggesting minimal multiple-cell bias during future downstream genomic processing from droplets

Loading an entire worm took <1 hour from droplet generation to FACS analysis, suggesting feasibility of collecting physiologically relevant data from droplet picoreactors. The successful cell loading of a whole complex organism demonstrates that the pipeline may have broad future applicability to other primary cell samples and diverse reaction schemes.

## CONCLUSION

In this work, we present a novel method capable of high-throughput phenotyping of large animal cells within double emulsion picoreactors compatible with FACS. Using our Dropception device and a commercial sorter, we encapsulated and screened 5 different mammalian cell lines (mouse: E14 ES, LM-1, NIH 3T3; human: Jurkat, HEK 293T) and primary cells dissociated fresh from a whole planarian flatworm in picoliter-scale droplets.

Double emulsions produced with our workflow are uniform, highly monodisperse, and stable under cell loading. In all cell lines, single-cell occupancy determined via microscopy approached maximal efficiency of ideal Poisson loading with no observed cell size bias. Importantly, droplet cell occupancies reported by high-throughput FACS mirror those calculated from microscopy, establishing that FACS can be used to screen cell-containing droplets at single-cell resolution. Encapsulation of primary cells from a whole planarian flatworm, a complex organism with high cellular diversity and technical challenges associated with sample preparation, performed as well as uniform cell lines with robust FACS phenotyping.

Unlike other droplet sorting strategies^41,42,77^, Dropception requires only commercially available equipment and limited technical expertise, making the technology easily adoptable by bioscience labs. For single-cell encapsulation, only an inexpensive microscope and syringe pumps are needed to operate the Dropception device; the setup is easy to use and can be assembled within a day. Downstream, we demonstrate high-throughput DE phenotyping using a widespread, inexpensive benchtop flow cytometer (Sony SH800). We have previously established that DE droplets can also be sorted using BD FACS Aria II and III machines^48^ and similarly anticipate Dropception will be compatible with most FACS instruments.

Furthermore, the Dropception pipeline reduces shear stresses on sorted cells via droplet shielding during FACS^78,79^, minimizing changes in underlying cell biology. The time required to load cells is short (< 30 minutes) and cells can be immediately lysed (and thus cell state ‘frozen’) upon encapsulation, enhancing the likelihood that recovered phenotypes accurately reflect native cell state^53^. Moreover, our unique device geometry allows for low flow rates and small droplet volumes, conserving samples and thus enabling new opportunities for single-cell analysis of rare or precious samples, including primary clinical samples, non-model organisms, or cell lines under chemical perturbation.

High-throughput screening and sorting of DE droplets containing complex animal cells opens up a wide range of potential applications. Sorting and sequencing only cell-containing droplets could vastly reduce reagent costs^15^ and increase accuracy for downstream next-generation sequencing, including eliminating common single-cell problems such as empty droplet false positives^32,33^. In addition, Dropception provides a method to screen cells based on a range of reaction- or secretion-based phenotypes traditionally inaccessible to FACS with enhanced sensitivity compared to other droplet techniques due to picoliter reaction volumes^80^.

In future work, we anticipate Dropception will facilitate a variety of multi-omic assays on the same single cell^16^ via isolation of single DE droplets for massively parallel downstream analysis. Using our prior sdDE-FACS pipeline for isolating individual DEs, single-cell droplets of a particular phenotype generated via the Dropception workflow could be sorted into wells of a multiwell plate for genome-wide processing, thereby diluting droplet buffers ~10,000-fold and thus enabling multiple assays per cell without a need for buffer exchange. Coupling genomic, epigenomic, or transcriptomic profiling to in-droplet cellular phenotyping with this scheme would allow direct investigation of genetic mechanisms driving cellular functions in the same cell using only simple plate-processing workflows. In this way, Dropception would expand the current repertoire of nucleic acid single-cell droplet assays to include functional, perturbation-based, or multi-omic profiling by coupling the throughput of droplet microfluidics with the power of flow cytometry.

## METHODS

### Extended Methods and Data Repository

Extended Methods are available in *Supplemental Information* including a step-by-step protocol for picoreactor single-cell encapsulation. Data, software, and the device design are available in an open-source Open Science Framework repository at: DOI 10.17605/OSF.IO/H4SR9.

### Cell Preparation and Viability Measurement

Cells were cultured according to ATCC standards. All cell lines, except LM1s, were dissociated using TrypLE Express Enzyme (Thermo Fisher Scientific). LM1s was dissociated using Accutase (STEMCELL Technologies). Planarian cells were collected by mechanical dissociation of whole animals using a razorblade and pipetting in 1X PBS. Prior to droplet loading, cells were stained with Calcein AM UltraBlue (AAT Bioquest), resuspended in 1X PBS + 0.04% (w/v) BSA, filtered with a 40 μm cell strainer, and diluted to a final calculated concentration of ~2.5 M/mL with 20% OptiPrep density gradient medium (Sigma) as projected from culture measurements (see *Extended Methods*). Viability and concentration measurements using a Trypan blue exclusion assay (**Table 2**) were conducted at multiple staging points using a Countess cell counter (Life Technologies).

### Double Emulsion Cell Encapsulation

Picoliter DEs were generated using four syringe pumps (PicoPump Elite, Harvard Apparatus) for cell suspension and inner, oil, and outer sheath solutions. Cell suspensions consisted of 1X PBS, 20% OptiPrep, 0.04% BSA, and diluted cells. The second inner phase was composed of 1X PBS with 0.5% BSA. The oil phase was composed of HFE7500 fluorinated oil (Sigma) and 2.2% Ionic PEG-Kyrtox^58^ (FSH, Miller-Stephenson). The carrier phase contained 1% Tween-20 (Sigma) and 2% Pluronix F68 (Kolliphor 188, Sigma) in PBS as described previously^48^. Each phase was loaded into syringes (PlastiPak, BD) and connected to the device via PE/2 tubing (Scientific Commodities). Typical flow rates were 400:125:105:6000 μl/hr (oil:cell:reagent:outer). A step-by-step protocol for cell preparation and device operation is available in *Extended Methods*.

### Picoreactor Phenotyping via Flow Cytometry

Single-cell DE picoreactors were analyzed via FACS using the sdDE-FACS workflow as described previously^48^. Briefly, 100 μL of collected droplets in 500 μL of sheath buffer containing 1% Tween-20 (Sigma) in 1X PBS were analyzed on the SH800 flow cytometer (Sony) using a standard 408 nm laser configuration and a 130 μm nozzle for sorting. After a brief pickup delay time (~2 min, as reported previously in the sdDE-FACS pipeline), DEs were gated on FSC-H *vs*. FSC-W for large particle analysis (population statistics, e.g., single core droplets *vs*. extraneous oil, confirmed via microscopy to identify gate) and subsequently gated on Calcein AM signal. All flow and thresholding parameters are reported in **Table S3**. Event rate was kept below 1,000 events/s for sorting purity.

## ASSOCIATED CONTENT

### Supporting Information

*Supplemental Information* including extended methods, a full workflow protocol, and referenced supplemental figures (SI, PDF). The device design, resources, and data are available via an Open Science Framework repository at: DOI 10.17605/OSF.IO/H4SR9.

## AUTHOR INFORMATION

### Author Contributions

‡These authors contributed equally. K.K.B., M.K., B.W. and P.M.F. conceived the study; K.K.B. and M.K. designed and directed the study. K.K.B. designed the method. P.H.S. and M.K. optimized workflow. K.K.B. and M.K. fabricated and characterized the device. K.K.B., M.K. and P.H.S. performed experiments. C.S. conducted cell culture and viability measurements. M.K. and G.K. contributed software. M.K. and K.K.B. conducted formal analysis, data curation, and validation. P.S., G.K., and S.G.K.C. conducted microscopy analysis and validation. S.Q. contributed resources. K.K.B., M.K., B.W., and P.M.F. wrote the manuscript, with input from all authors. All authors have given approval to the final version of the manuscript.

### Notes

Methods and techniques outlined in this work are disclosed in a U.S. patent filing, U.S. PTO Application No. 62/693,800, filed by co-authors K.K.B and P.M.F.

## ACKNOWLEDGMENTS

The authors acknowledge members of the Fordyce and Wang laboratories for their feedback, assistance, and suggestions in review of the manuscript. In particular, we acknowledge Conor McClune for his assistance in droplet counting and critical feedback, as well as the larger Dropception team for supporting this work. We acknowledge Dr. David Sukovich and Dr. Adam Abate (UCSF) for sharing their flow-focusing device design, which was modified to include new design elements in this study. We additionally thank the Stanford FACS Core, Stanford Protein and Nucleic Acid (PAN) and Stanford Nano Shared Facilities (SNSF), supported by the National Science Foundation under award ECCS-1542152, for use of their instruments and facilities. This work was supported by a NIH grant 1DP2GM123641 (P.M.F.) and a HFSP grant RGY0085/2019 (B.W.). K.K.B. acknowledges support as Chem-H CBI fellow (NIHT32 GM 120007), an NSF GFRP fellow, and a Siebel Scholar. M.K. acknowledges support as a BioX SIGF Lavidge and McKinley fellow. P.M.F. is a Chan Zuckerberg Biohub Investigator.

## ABBREVIATIONS

FACS: fluorescence-activated cell sorting
DE: double emulsion

## References

(1) Svensson, V.; Vento-Tormo, R.; Teichmann, S. A. Exponential Scaling of Single-Cell RNA-Seq in the Past Decade. Nat Protoc 2018, 13, 599–604.

(2) Yuan, G.-C.; Cai, L.; Elowitz, M.; Enver, T.; Fan, G.; Guo, G.; Irizarry, R.; Kharchenko, P.; Kim, J.; Orkin, S.; Quackenbush, J.; Saadatpour, A.; Schroeder, T.; Shivdasani, R.; Tirosh, I. Challenges and Emerging Directions in Single-Cell Analysis. Genome Biol 2017, 18, 84.

(3) Macosko, E. Z.; Basu, A.; Satija, R.; Nemesh, J.; Shekhar, K.; Goldman, M.; Tirosh, I.; Bialas, A. R.; Kamitaki, N.; Martersteck, E. M.; Trombetta, J. J.; Weitz, D. A.; Sanes, J. R.; Shalek, A. K.; Regev, A.; McCarroll, S. A. Highly Parallel Genome-Wide Expression Profiling of Individual Cells Using Nanoliter Droplets. Cell 2015, 161 (5), 1202–1214.

(4) Klein, A. M.; Mazutis, L.; Akartuna, I.; Tallapragada, N.; Veres, A.; Li, V.; Peshkin, L.; Weitz, D. A.; Kirschner, M. W. Droplet Barcoding for Single-Cell Transcriptomics Applied to Embryonic Stem Cells. Cell 2015, 161 (5), 1187–1201.

(5) Buenrostro, J. D; Wu, B.; Litzenburger, U. M.; Ruff, D.; Gonzales, M. L.; Snyder, M. P.; Chang, H. Y.; Greenleaf, W. J. Single-Cell Chromatin Accessibility Reveals Principles of Regulatory Variation. Nature 2015, 523, 486–490.

(6) Rotem, A; Ram, O.; Shoresh, N.; Sperling, R. A.; Goren, A.; Weitz, D. A.; Bernstein, B. E. Single-Cell ChIP-Seq Reveals Cell Subpopulations Defined by Chromatin State. Nat Biotechnol 2015, 33, 1165–1172.

(7) Nagano, T.; Lubling, Y.; Stevens, T. J.; Schoenfelder, S.; Yaffe, E.; Dean, W.; Laue, E. D.; Tanay, A.; Fraser, P. Single-Cell Hi-C Reveals Cell-to-Cell Variability in Chromosome Structure. Nature 2013, 502, 59–64.

(8) Achim, K.; Pettit, J.-B.; Saraiva, L. R.; Gavriouchkina, D.; Larsson, T.; Arendt, D.; Marioni, J. C. High-Throughput Spatial Mapping of Single-Cell RNA-Seq Data to Tissue of Origin. Nat Biotechnol 2015, 33, 503–509.

(9) Lee, J. H.; Daugharthy, E. R.; Scheiman, J.; Kalhor, R.; Yang, J. L.; Ferrante, T. C.; Terry, R.; Jeanty, S. S. F.; Li, C.; Amamoto, R.; Peters, D. T.; Turczyk, B. M.; Marblestone, A. H.; Inverso, S. A.; Bernard, A.; Mali, P.; Rios, X.; Aach, J.; Church, G. M. Highly Multiplexed Subcellular RNA Sequencing in Situ. Science 2014, 343 (6177), 1360–1363.

(10) Battich, N.; Stoeger, T.; Pelkmans, L. Image-Based Transcriptomics in Thousands of Single Human Cells at Single-Molecule Resolution. Nat Methods 2013, 10, 1127–1133.

(11) Schmidt, S. T.; Zimmerman, S. M.; Wang, J.; Kim, S. K.; Quake, S. R. Quantitative Analysis of Synthetic Cell Lineage Tracing Using Nuclease Barcoding. ACS Synth Biol 2017, 6 (6), 936–942.

(12) Trapnell, C.; Cacchiarelli, D.; Grimsby, J.; Pokharel, P.; Li, S.; Morse, M.; Lennon, N. J.; Livak, K. J; Mikkelsen, T. S.; Rinn, J. L. The Dynamics and Regulators of Cell Fate Decisions Are Revealed by Pseudotemporal Ordering of Single Cells. Nat Biotechnol 2014, 32, 381–386.

(13) Alemany, A.; Florescu, M.; Baron, C. S.; Peterson-Maduro, J.; Oudenaarden, A. Whole-Organism Clone Tracing Using Single-Cell Sequencing. Nature 2018, 556, 108–112.

(14) McKenna, A.; Findlay, G. M.; Gagnon, J. A.; Horwitz, M. S.; Schier, A. F.; Shendure, J. Whole-Organism Lineage Tracing by Combinatorial and Cumulative Genome Editing. Science 2016, 353 (6298), aaf7907.

(15) Zhang, X.; Li, T.; Liu, F.; Chen, Y.; Yao, J.; Li, Z.; Huang, Y.; Wang, J. Comparative Analysis of Droplet-Based Ultra-High-Throughput Single-Cell RNA-Seq Systems. Mol Cell 2019, 73 (1), 130–142.e5.

(16) Stuart, T.; Satija, R. Integrative Single-Cell Analysis. Nat Rev Gen 2019, 20, 257–272.

(17) Wen, N.; Zhao, Z.; Fan, B.; Chen, D.; Men, D.; Wang, J.; Chen, J. Development of Droplet Microfluidics Enabling High-Throughput Single-Cell Analysis. Molecules 2016, 21 (7), 881.

(18) Fu, Y.; Li, C.; Lu, S.; Zhou, W.; Tang, F.; Xie, X. S.; Huang, Y. Uniform and Accurate Single-Cell Sequencing Based on Emulsion Whole-Genome Amplification. Proc Natl Acad Sci 2015, 112 (38), 11923–11928.

(19) Lan, F.; Demaree, B.; Ahmed, N.; Abate, A. R. Single-Cell Genome Sequencing at Ultra-High Throughput with Microfluidic Droplet Barcoding. Nat Biotechnol 2017, 35, 640–646.

(20) Satpathy, A. T.; Granja, J. M.; Yost, K. E.; Qi, Y.; Meschi, F.; McDermott, G. P.; Olsen, B. N.; Mumbach, M. R.; Pierce, S. E.; Corces, M. R.; Shah, P.; Bell, J. C.; Jhutty, D.; Nemec, C. M.; Wang, J.; Wang, L.; Yin, Y.; Giresi, P. G.; Chang, A. L. S.; Zheng, G. X. Y.; Greenleaf, W. J.; Chang, H. Y. Massively Parallel Single-Cell Chromatin Landscapes of Human Immune Cell Development and Intratumoral T Cell Exhaustion. Nat Biotechnol 2019, 37, 925–936.

(21) Lareau, C. A.; Duarte, F. M.; Chew, J. G.; Kartha, V. K.; Burkett, Z. D.; Kohlway, A. S.; Pokholok, D.; Aryee, M. J.; Steemers, F. J.; Lebofsky, R.; Buenrostro, J. D. Droplet-Based Combinatorial Indexing for Massive-Scale Single-Cell Chromatin Accessibility. Nat Biotechnol 2019, 37, 916–924.

(22) Zheng, G. X. Y.; Terry, J. M.; Belgrader, P.; Ryvkin, P.; Bent, Z. W.; Wilson, R.; Ziraldo, S. B.; Wheeler, T. D.; McDermott, G. P.; Zhu, J.; Gregory, M. T.; Shuga, J.; Montesclaros, L.; Underwood, J. G.; Masquelier, D. A.; Nishimura, S. Y.; Schnall-Levin, M.; Wyatt, P. W.; Hindson, C. M.; Bharadwaj, R.; Wong, A.; Ness, K. D.; Beppu, L. W.; Deeg, H. J.; McFarland, C.; Loeb, K. R.; Valente, W. J.; Ericson, N. G.; Stevens, E. A.; Radich, J. P.; Mikkelsen, T. S.; Hindson, B. J.; Bielas, J. H.. Massively Parallel Digital Transcriptional Profiling of Single Cells. Nat Commun 2017, 8, 14049.

(23) Shahi, P.; Kim, S. C.; Haliburton, J. R.; Gartner, Z. J.; Abate, A. R. Abseq: Ultrahigh-Throughput Single Cell Protein Profiling with Droplet Microfluidic Barcoding. Sci Rep 2017, 7, 44447.

(24) Stoeckius, M.; Hafemeister, C.; Stephenson, W.; Houck-Loomis, B.; Chattopadhyay, P. K.; Swerdlow, H.; Satija, R.; Smibert, P. Simultaneous Epitope and Transcriptome Measurement in Single Cells. Nat Methods 2017, 14, 865–868.

(25) Mimitou, E. P.; Cheng, A.; Montalbano, A.; Hao, S.; Stoeckius, M.; Legut, M.; Roush, T.; Herrera, A.; Papalexi, E.; Ouyang, Z.; Satija, R.; Sanjana, N. E.; Koralov, S. B.; Smibert, P. Multiplexed Detection of Proteins, Transcriptomes, Clonotypes and CRISPR Perturbations in Single Cells. Nat Methods 2019, 16, 409–412.

(26) Karaiskos, N.; Wahle, P.; Alles, J.; Boltengagen, A.; Ayoub, S.; Kipar, C.; Kocks, C.; Rajewsky, N.; Zinzen, R. P. The Drosophila Embryo at Single-Cell Transcriptome Resolution. Science 2017, 358 (6360), 194–199.

(27) The Tabula Muris Consortium. Single-Cell Transcriptomics of 20 Mouse Organs Creates a Tabula Muris. Nature 2018, 562 (7727), 367–372.

(28) Wagner, D. E.; Weinreb, C.; Collins, Z. M.; Briggs, J. A.; Megason, S. G.; Klein, A. M. Single-Cell Mapping of Gene Expression Landscapes and Lineage in the Zebrafish Embryo. Science 2018, 360 (6392), 981–987.

(29) Fincher, C. T.; Wurtzel, O.; de Hoog, T.; Kravarik, K. M.; Reddien, P. W. Cell Type Transcriptome Atlas for the Planarian *Schmidtea Mediterranea*. Science 2018, 360 (6391), eaaq1736.

(30) Suea-Ngam, A.; Howes, P. D.; Srisa-Art, M.; deMello, A. J. Droplet Microfluidics: From Proof-of-Concept to Real-World Utility? Chem Commun 2019, 55 (67), 9895–9903.

(31) DePasquale, E. A. K.; Schnell, D. J.; Van Camp, P.-J.; Valiente-Alandí, Í.; Blaxall, B. C.; Grimes, H. L.; Singh, H.; Salomonis, N. DoubletDecon: Deconvoluting Doublets from Single-Cell RNA-Sequencing Data. Cell Reports 2019, 29 (6), 1718–1727.e8.

(32) Petukhov, V.; Guo, J.; Baryawno, N.; Severe, N.; Scadden, D. T.; Samsonova, M. G.; Kharchenko, P. V. DropEst: Pipeline for Accurate Estimation of Molecular Counts in Droplet-Based Single-Cell RNA-Seq Experiments. Genome Biol 2018, 19, 78.

(33) Lun, A. T. L.; Riesenfeld, S.; Andrews, T.; Dao, T. P.; Gomes, T.; Marioni, J. C. EmptyDrops: Distinguishing Cells from Empty Droplets in Droplet-Based Single-Cell RNA Sequencing Data. Genome Biol 2019, 20, 63

(34) Herzenberg, L. A.; Parks, D.; Sahaf, B.; Perez, O.; Roederer, M.; Herzenberg, L. A. The History and Future of the Fluorescence Activated Cell Sorter and Flow Cytometry: A View from Stanford. Clin Chem 2002, 48 (10), 1819–1827.

(35) Chattopadhyay, P. K.; Roederer, M. A Mine Is a Terrible Thing to Waste: High Content, Single Cell Technologies for Comprehensive Immune Analysis: Single Cell Technologies for Immune Analysis. Am J Transplant 2015, 15 (5), 1155–1161.

(36) Dhar, M.; Lam, J. N.; Walser, T.; Dubinett, S. M.; Rettig, M. B.; Di Carlo, D. Functional Profiling of Circulating Tumor Cells with an Integrated Vortex Capture and Single-Cell Protease Activity Assay. Proc Natl Acad Sci 2018, 115 (40), 9986–9991.

(37) Chokkalingam, V.; Tel, J.; Wimmers, F.; Liu, X.; Semenov, S.; Thiele, J.; Figdor, C. G.; Huck, W. T. S. Probing Cellular Heterogeneity in Cytokine-Secreting Immune Cells Using Droplet-Based Microfluidics. Lab Chip 2013, 13 (24), 4740–4744.

(38) Eyer, K.; Doineau, R. C. L.; Castrillon, C. E.; Briseño-Roa, L.; Menrath, V.; Mottet, G.; England, P.; Godina, A.; Brient-Litzler, E.; Nizak, C.; Jensen, A.; Griffiths, A. D.; Bibette, J.; Bruhns, P.; Baudry, J. Single-Cell Deep Phenotyping of IgG-Secreting Cells for High-Resolution Immune Monitoring. Nat Biotechnol 2017, 35, 977–982.

(39) Aharoni, A.; Amitai, G.; Bernath, K.; Magdassi, S.; Tawfik, D. S. High-Throughput Screening of Enzyme Libraries: Thiolactonases Evolved by Fluorescence-Activated Sorting of Single Cells in Emulsion Compartments. Chem Biol 2005, 12 (12), 1281–1289.

(40) Mastrobattista, E.; Taly, V.; Chanudet, E.; Treacy, P.; Kelly, B. T.; Griffiths, A. D. High-Throughput Screening of Enzyme Libraries: In Vitro Evolution of a β-Galactosidase by Fluorescence-Activated Sorting of Double Emulsions. Chem Biol 2005, 12 (12), 1291–1300.

(41) Baret, J.-C.; Miller, O. J.; Taly, V.; Ryckelynck, M.; El-Harrak, A.; Frenz, L.; Rick, C.; Samuels, M. L.; Hutchison, J. B.; Agresti, J. J.; Link, D. R.; Weitz, D. A.; Griffiths, A. D. Fluorescence-Activated Droplet Sorting (FADS): Efficient Microfluidic Cell Sorting Based on Enzymatic Activity. Lab Chip 2009, 9 (13), 1850–1858.

(42) Cole, R. H.; Tang, S.-Y.; Siltanen, C. A.; Shahi, P.; Zhang, J. Q.; Poust, S.; Gartner, Z. J.; Abate, A. R. Printed Droplet Microfluidics for on Demand Dispensing of Picoliter Droplets and Cells. Proc Natl Acad Sci 2017, 114 (33), 8728–8733.

(43) Mazutis, L.; Gilbert, J.; Ung, W. L.; Weitz, D. A.; Griffiths, A. D.; Heyman, J. A. Single-Cell Analysis and Sorting Using Droplet-Based Microfluidics. Nat Protoc 2013, 8, 870–891.

(44) Zinchenko, A.; Devenish, S. R. A.; Kintses, B.; Colin, P.-Y.; Fischlechner, M.; Hollfelder, F. One in a Million: Flow Cytometric Sorting of Single Cell-Lysate Assays in Monodisperse Picolitre Double Emulsion Droplets for Directed Evolution. Anal Chem 2014, 86 (5), 2526–2533.

(45) Shapiro, H. M. Practical Flow Cytometry, 4th ed.; Wiley-Liss: New York, 2003.

(46) Tu, R.; Martinez, R.; Prodanovic, R.; Klein, M.; Schwaneberg, U. A Flow Cytometry–Based Screening System for Directed Evolution of Proteases. J Biomol Screen 2011, 16 (3), 285–294.

(47) Ma, F.; Xie, Y.; Huang, C.; Feng, Y.; Yang, G. An Improved Single Cell Ultrahigh Throughput Screening Method Based on In Vitro Compartmentalization. PLoS ONE 2014, 9 (2), e89785.

(48) Brower, K. K.; Carswell-Crumpton, C.; Klemm, S.; Cruz, B.; Kim, G.; Calhoun, S. G. K.; Nichols, L.; Fordyce, P. M. Double Emulsion Flow Cytometry with High-Throughput Single Droplet Isolation and Nucleic Acid Recovery. Lab Chip 2020, 10.1039.D0LC00261E.

(49) Collins, D. J.; Neild, A.; deMello, A.; Liu, A.-Q.; Ai, Y. The Poisson Distribution and beyond: Methods for Microfluidic Droplet Production and Single Cell Encapsulation. Lab Chip 2015, 15 (17), 3439–3459.

(50) Hümmer, D.; Kurth, F.; Naredi-Rainer, N.; Dittrich, P. S. Single Cells in Confined Volumes: Microchambers and Microdroplets. Lab Chip 2016, 16 (3), 447–458.

(51) Lim, S. W.; Abate, A. R. Ultrahigh-Throughput Sorting of Microfluidic Drops with Flow Cytometry. Lab Chip 2013, 13 (23), 4563–4572.

(52) Sukovich, D. J.; Lance, S. T.; Abate, A. R. Sequence Specific Sorting of DNA Molecules with FACS Using 3dPCR. Sci Rep 2017, 7, 39385.

(53) Nguyen, Q. H.; Pervolarakis, N.; Nee, K.; Kessenbrock, K. Experimental Considerations for Single-Cell RNA Sequencing Approaches. Front Cell Dev Biol 2018, 6, 108.

(54) Silva, B. F. B.; Rodríguez-Abreu, C.; Vilanova, N. Recent Advances in Multiple Emulsions and Their Application as Templates. Curr Opin Colloid Interface Sci 2016, 25, 98–108.

(55) Chong, D.; Liu, X.; Ma, H.; Huang, G.; Han, Y. L.; Cui, X.; Yan, J.; Xu, F. Advances in Fabricating Double-Emulsion Droplets and Their Biomedical Applications. Microfluid Nanofluid 2015, 19, 1071–1090.

(56) Abate, A. R.; Thiele, J.; Weitz, D. A. One-Step Formation of Multiple Emulsions in Microfluidics. Lab Chip 2011, 11 (2), 253–258.

(57) Kim, S. C.; Sukovich, D. J.; Abate, A. R. Patterning Microfluidic Device Wettability with Spatially-Controlled Plasma Oxidation. Lab Chip 2015, 15 (15), 3163–3169.

(58) Holtze, C.; Rowat, A. C.; Agresti, J. J.; Hutchison, J. B.; Angilè, F. E.; Schmitz, C. H. J.; Köster, S.; Duan, H.; Humphry, K. J.; Scanga, R. A.; Johnson, J. S.; Pisignano, D.; Weitz, D. A. Biocompatible Surfactants for Water-in-Fluorocarbon Emulsions. Lab Chip 2008, 8 (10), 1632–1639.

(59) Chen, F.; Zhan, Y.; Geng, T.; Lian, H.; Xu, P.; Lu, C. Chemical Transfection of Cells in Picoliter Aqueous Droplets in Fluorocarbon Oil. Anal Chem 2011, 83 (22), 8816–8820.

(60) Edd, J. F.; Di Carlo, D.; Humphry, K. J.; Köster, S.; Irimia, D.; Weitz, D. A.; Toner, M. Controlled Encapsulation of Single-Cells into Monodisperse Picolitre Drops. Lab Chip 2008, 8 (8), 1262–1264.

(61) Chabert, M.; Viovy, J.-L. Microfluidic High-Throughput Encapsulation and Hydrodynamic Self-Sorting of Single Cells. Proc Nat Acad Sci 2008, 105 (9), 3191–3196.

(62) Gerver, R. E.; Gómez-Sjöberg, R.; Baxter, B. C.; Thorn, K. S.; Fordyce, P. M.; Diaz-Botia, C. A.; Helms, B. A.; DeRisi, J. L. Programmable Microfluidic Synthesis of Spectrally Encoded Microspheres. Lab Chip 2012, 12 (22), 4716–4723.

(63) Jiang, N.; Hu, Y.; Liu, X.; Wu, Y.; Zhang, H.; Chen, G.; Liang, J.; Lu, X.; Liu, S. Differentiation of E14 Mouse Embryonic Stem Cells into Thyrocytes *In Vitro*. Thyroid 2010, 20 (1), 77–84.

(64) Incarnato, D.; Neri, F. High-Throughput Whole-Genome Sequencing of E14 Mouse Embryonic Stem Cells. Genom Data 2015, 3, 6–7.

(65) Liu, Y.; Yu, C.; Daley, T. P.; Wang, F.; Cao, W. S.; Bhate, S.; Lin, X.; Still, C.; Liu, H.; Zhao, D.; Wang, H.; Xie, X. S.; Ding, S.; Wong, W. H.; Wernig, M.; Qi, L. S. CRISPR Activation Screens Systematically Identify Factors That Drive Neuronal Fate and Reprogramming. Cell Stem Cell 2018, 23 (5), 758–771.e8.

(66) Abraham, R. T.; Weiss, A. Jurkat T Cells and Development of the T-Cell Receptor Signalling Paradigm. Nat Rev Immunol 2004, 4, 301–308.

(67) Chi, S.; Weiss, A.; Wang, H. A CRISPR-Based Toolbox for Studying T Cell Signal Transduction. Biomed Res Int 2016, 2016, 5052369.

(68) Kim, H.; Bojar, D.; Fussenegger, M. A CRISPR/Cas9-Based Central Processing Unit to Program Complex Logic Computation in Human Cells. Proc Natl Acad Sci 2019, 116 (15), 7214–7219.

(69) Montano, M. Model Systems. In Translational Biology in Medicine; Elsevier, 2014; pp 9–33.

(70) Shang, L.; Cheng, Y.; Zhao, Y. Emerging Droplet Microfluidics. Chem Rev 2017, 117 (12), 7964–8040.

(71) Newmark, P. A.; Alvarado, A. S. Not Your Father#8217;s Planarian: A Classic Model Enters the Era of Functional Genomics. Nat Rev Genet 2002, 3, 210–219.

(72) Reddien, P. W.; Alvarado, A. S. Fundamentals of Planarian Regeneration. Annu Rev Cell Dev Biol 2004, 20, 725–757.

(73) Wagner, D. E.; Wang, I. E.; Reddien, P. W. Clonogenic Neoblasts Are Pluripotent Adult Stem Cells That Underlie Planarian Regeneration. Science 2011, 332 (6031), 811–816.

(74) Rink, J. C. Stem Cell Systems and Regeneration in Planaria. Dev Genes Evol 2013, 223 (1–2), 67–84.

(75) Reddien, P. W.; Bermange, A. L.; Murfitt, K. J.; Jennings, J. R.; Sánchez Alvarado, A. Identification of Genes Needed for Regeneration, Stem Cell Function, and Tissue Homeostasis by Systematic Gene Perturbation in Planaria. Dev Cell 2005, 8 (5), 635–649.

(76) Alvarado, A. S.; Tsonis, P. A. Bridging the Regeneration Gap: Genetic Insights from Diverse Animal Models. Nat Rev Genet 2006, 7 (11), 873–884.

(77) Eastburn, D. J.; Huang, Y.; Pellegrino, M.; Sciambi, A.; Ptáček, L. J.; Abate, A. R. Microfluidic Droplet Enrichment for Targeted Sequencing. Nucleic Acids Res 2015, 43 (13), e86–e86.

(78) Chen, Y.; Liu, X.; Zhao, Y. Deformation Dynamics of Double Emulsion Droplet under Shear. Appl Phys Lett 2015, 106 (14), 141601.

(79) Chen, Y.; Liu, X.; Zhang, C.; Zhao, Y. Enhancing and Suppressing Effects of an Inner Droplet on Deformation of a Double Emulsion Droplet under Shear. Lab on a Chip 2015, 15 (5), 1255–1261.

(80) Streets, A. M.; Huang, Y. Microfluidics for Biological Measurements with Single-Molecule Resolution. Curr Opin Biotechnol 2014, 25, 69–77.

